# Mice follow odor trails using stereo olfactory cues and rapid sniff to sniff comparisons

**DOI:** 10.1101/293746

**Authors:** Peter W Jones, Nathan N Urban

**Author notes:** Corresponding author: Nathan N Urban Department of Neurobiology, Center for the Neural Basis of Cognition, University of Pittsburgh 3500 Terrace Street, STE E1446 Pittsburgh, PA 15213-2500.

## Abstract

Animals use the distribution of chemicals in their environment to guide behaviors essential for life, including finding food and mates, and avoiding predators. The presence of this general class of behavior is extremely widespread, even though the olfactory sensory apparatus and strategies used may vary between animals. The strategies and cues used by mammals to localize and track odor sources have recently become of interest to neuroscientists, but are still poorly understand. In order to study how mice localize odors, we trained mice to perform a trail following task using a novel behavioral paradigm and behavioral monitoring setup. We find that mice, in order to follow an odor trail, use both sniff by sniff odor concentration comparisons and internares comparisons to guide their behavior. Furthermore, they employ olfactory information to guide adaptive behaviors with remarkably short latencies of approximately 80ms. This study and its findings establish a rich, quantifiable olfactory localization behavior in mice that is amenable to physiological investigations and motivates investigation into the neural substrates of the identified olfactory cues.

**Significance:** Many animals, like rodents, rely heavily on their sense smell to guide them as they navigate their environment, to find food and mates and to avoid predators. Yet, in mammals, this function of the olfactory system is much less well studied than odor identification. We created a behavioral task where mice had to follow odor trails in order to efficiently find food and then tracked their movements around those trails. We found that they respond to sniff-by-sniff changes in odor intensity, using those changes to guide movements in less than one tenth of a second and also confirm that they use stereo cues to guide behavior. These results lay the groundwork for determining the brain circuits underlying olfactory navigation.

## Introduction

Across the animal kingdom, animals use their olfactory systems to localize odor sources for a variety of purposes vital for the animal’s survival and viability, including finding food sources and mates, following pheromone trails, and avoiding predators. In all of these cases, the animal must orient and move based on samples of the chemical environment taken over time and often using multiple sensors (e.g. paired nares or antennae). Detailed behavioral and modeling studies find that animals use a variety of strategies to navigate chemical gradients. Bacteria and nematode worms seem to ascend attractant odorant gradients (chemotaxis) using biased random walks (Pierce-Shimomura et al., 1999). Some species may use multisensory strategies to localize sources (Collett and Cardé, 2014; McMeniman et al., 2014). Others employ strategies involving consistent reorientation toward the increasing axis of the gradient (Porter et al., 2007; Louis et al., 2008; Khan et al., 2012; Perna et al., 2012; Catania, 2013).

Most of the detailed work describing strategies by which terrestrial animals use olfactory cues to guide their movements has been performed using invertebrate models and computational modeling based on those results. Analysis of ant trail following while foraging has led to several detailed models of how ants follow local olfactory cues to stay along the trial (Calenbuhr and Deneubourg, 1992; Perna et al., 2012). These models successfully describe ant trail following behavior using the simultaneous concentrations of odorant detected by each of two antennae. However, in the absence of the ability to compare signals from the two antennae, such as when one is removed, ants can still follow pheromone trails, though less well (Hangartner, 1967). The ability to follow odor gradients with only one sensor has similarly been demonstrated in bees and fly larvae (Martin, 1965; Louis et al., 2008). When only one olfactory sensor is present, the animal must rely on self-movement and comparisons of olfactory concentrations over time in order to estimate the local odor gradient, which may be facilitated by a periodic, side-to-side movement termed “casting” (Kennedy, 1983).

The mechanisms vertebrates use to localize and follow odor sources are poorly understood, even though some mammals show remarkable olfactory tracking capabilities, e.g. (Thesen et al., 1993). Experiments in several mammals, including humans, have shown that stereo (inter-nares) olfactory cues are used to orient towards odor sources and follow trails (Porter et al., 2007; Khan et al., 2012; Catania, 2013; Rabell et al., 2017). Those results also demonstrate that mammals can localize and track odors without stereo cues; they consistently show impaired performance when a single nares is occluded, but not total inability to perform the task. However, the precise cues and strategies guiding mammalian odor tracking behavior are largely unknown.

Understanding what cues are being used by animals for olfactory source localization and how those cues are being used to guide movement are important first steps in studying the neural processes governing such behavior. Towards this end, we trained mice to follow surface-applied odor trials and quantified their behavior in detail. We find that mice use primarily sniff-to-sniff comparisons of olfactory input in order to make decisions about changes of direction. The mice use this olfactory information remarkably quickly, with turning following sniffing by less than 80 ms. We additionally identify that, in this behavior, stereo (inter-nares) olfactory cues impair precise localization of the trail on the millimeter scale.

## Materials and Methods

### Behavioral Arena

A custom behavioral arena was designed and constructed to provide evenly-illuminated, glare and obstruction free video monitoring of the mouse’s nose position over a its large surface. The floor of the behavioral arena measures 36×45” and is made up of a stack of materials (bottom to top): 3/8” Starphire low-iron glass (PPG), 10mm Endlighten acrylic (Evonik), and thin abrasion resistant acrylic. Infrared (IR) LEDs (www.environmentallights.com) provided the illumination for video monitoring and were mounted into a polished aluminum U-channel frame to edge-on illuminate the Endlighten acrylic layer. The floor was then mounted into an aluminum rail table (80/20 Inc) designed to hold the floor. The sides were left open; though mice do peer out over the edge, especially during initial behavioral sessions, they did not jump or run off the edge.

A machine vision camera with a native resolution for 1280×1024 @ 150fps was mounted under the arena floor to record behavior (Flea3, Point Grey Imaging; Lens – Computar 5mm). We captured movies at 30-60fps for this study, with a linear resolution of .832 mm/px. The primary reason for taking video from below is so that the animal’s nose can be continuously tracked while it is investigating the surface. Initial attempts at monitoring tracking from directly above resulted in many instances when the nose was occluded from view by other body parts. During behavior, all ambient light sources in the room (computer monitors, etc) were deep red filtered at wavelengths longer than visual spectrum of the mouse.

### Animal training procedures

Mice used in this study were C57/BI6 genetic background, and were of both sexes, but predominantly female. The mice were food restricted, in accordance with Carnegie Mellon University Animal Care and Use guidelines, in order to ensure motivation for seeking food rewards during behavior. Mice performed 3-8 odor tracking trials per day for food reward. On each trial, mice were removed from their cage, via a transfer tube, and placed onto the edge of the arena, near the edge of an odor trail. The rewarded trail was baited with small bits of food (peanut or chocolate) intermittently along its length, the inter-reward distance was variable during the course of training, up to ˜80 cm, and was gradually increased as the mice became more willing to follow the odor trail. Trail end points, overall locations, and shapes are varied trial to trial to be unpredictable. Trail complexity was also increased during the course of training; trails were initially short and straight, and gradually increased in length and complexity to span the table width while containing significant curves, corners, and crossings between the two trails. We did this to convince ourselves that the mice were really following the trail and not merely investigating along an approximately straight path. An animal saw each odor trail at most twice, and if a trail was used twice, the animal was released at opposite ends of the trail so that it never was expected to follow the same path twice.

Training on this task took approximately 16 daily behavioral sessions. The mice were allowed two sessions, 5 minutes each, to habituate to the arena before exposure to odor trails. Food was randomly scattered on the table surface for these sessions. Mice generally spent the majority of their time around the edges of the arena, occasionally crossing the center, during the first session. Most mice began eating the food during the second arena exposure session. On the 3rd session, short trails were drawn and food was only place on the rewarded trail. The first rewarded trails were almost entirely covered with reward, which were rapidly decreased in density as the trails were drawn with increased length and complexity during the training period. Trained mice regularly followed trails of >1.2m length, with sections of >40 cm in length where no food was present.

### Nares occlusion

Mice were nares occluded under ketamine/xylezene anesthesia using reversible plugs, following Cummings et al (1997). After insertion, a successfully occluding seal was indicated by the absence of drainage or bubbles after applying soapy water to the nares opening. Occlusions were re-checked halfway through each occlusion period and upon removal. Data collected before any failed occlusion checks were excluded from the analysis.

### Respiration Monitoring

Air temperature changes during respiration were recorded using a 40 gauge, PFA insulated, T-type thermocouple (Omega) implanted in the left nasal cavity of a subset (n=5) of mice. During surgery, mice were anesthetized using isofluorane, and analgesia administered both pre and post surgically according to Animal Care and Use Guidelines. A burr hole was drilled through the bone above the nasal cavity, and a syringe needle was used to pierce the underlying tissue. The thermocouple was lowered into the cavity using a micromanipulator while its signal was monitored. The precise placement in depth was judged by the experimenter. Sometimes, cues to guide placement, such as periodic temperature changes or a step increase in temperature corresponding to the probe touching the bottom of the cavity, could be observed. A head restraint bar was affixed to the skull during the same surgery so that animals could be temporarily restrained in order to connect leads each day. Both the thermocouple and head bar were fixed to the skull using dental acrylic. Animals were given 3-4 days to recover before restarting behavior.

Recorded respiration signals were low pass filtered at l-2kHz and amplified by an analog amplifier (Brown-Lee), and digitized at 5kHz by a 16bit D/A interface (Instrutech). Signals were digitally filtered with a 5ms width median filter, and baseline fluctuations were removed through subtraction of a low pass filtered version of the signal produced by convolution with a Gaussian filter (σ= 40 ms). The inspiration peaks were then detected in this high-pass filtered respiration signal, and the resulting binary vector was smoothed by convolving with a Gaussian kernel (σ=50 ms) to yield the continuous estimate of sniff rate. Using inter-sniff interval based sniff rates yields qualitatively similar results to those reported here (not shown).

### Automated tracking and behavioral analysis

Automated tracking was performed using saved video of behavior. If respiration data were also collected, video frame acquisition was triggered by the data acquisition software, Igor Pro (Wavemetrics), in order to synchronize video with the thermocouple signal. All data and statistical analyses were performed in MATLAB (Mathworks) using built-in functionality and custom programs.

Automated, post-hoc video tracking was performed on the period of time from which the mice were first put on the table until the time at which they completed following the full length of the trail or they no longer paid attention to the trail, approximately 30 seconds to 1.5 minutes for well-trained animals. During early trials, the animals regularly explored other regions of the environment between bouts of trail exploration, thus the animals would be given up to 5 minutes to explore the trail. For calculation of behavioral measures such as ‘following efficiency’ (defined in Results), the entire length of analyzed video was used as the time period of the analysis.

The automated tracking algorithm was implemented using standard computer vision techniques and will be made available for use per request to the authors. Briefly, the program subtracts an average frame to isolate the moving portions of the scene (the mouse), then thresholds that image to make a binary image of detected “blobs” and background. The threshold is set at a level so that the tracked blobs are the animal’s nose, paws, and tail. In every frame, each blob is judged to be new or previously identified based on its positional similarity to blobs in the last frame, and is given a unique ID if new. Blob labeling as tail or nose was performed heuristically on each frame as follows. The tail was first identified based on its elongated aspect ratio. For a mouse with straight body posture, the nose is expected to be the blob that is farthest along the body axis from the tail. Thus the blob furthest along the body axis from the tail is labeled as ‘nose’ if it is farther than a threshold distance, which was set conservatively to exclude paws being falsely labeled as ‘nose’ during frames when the animal was in a bent posture. This algorithm leaves the ‘nose’ unidentified in some frames, therefor labels are propagated to blobs in adjacent frames with the same ID, allowing the nose, which is easily identifiable from paws when the animal is stretched out, to be identified in frames when the animal is bent into postures where nose and paw are difficult to distinguish based solely on their size and position in that single frame. The tracking output of all videos was subsequently reviewed for accurate tracking and manually corrected if necessary. All nose positions in this manuscript are the centroid of the blob identified as the nose by this procedure.

The scented wax trails were bright appearing on video. Their contours were semi-automatically detected using canny edge detection and filling operations (Canny, 1986; Szeliski, 2010). The trail detection algorithm runs after it is first seeded with a point by clicking on the trail in a graphical user interface. The full detected trail was then reviewed by the experimenter for verification and removal of any spuriously detected trail segments.

### Analysis of nose positions during tracking

We analyzed sections of behavior when the animal was within 20 px (16.68mm) of the trail (left is negative, right is positive), which corresponded to the approximate distance within which the mouse appeared to actively be engaged in searching along the trail. For analysis, segments of following behavior are then concatenated together, with no segment shorter than 40 position samples, to generate a single time-series of following behavior that spans all behavioral sessions for a single mouse.

### Experimental Design and Statistical Analysis

Nonparameteric statistical tests, e.g. Wilcoxon Rank-Sum, are used throughout this study to compare the medians of measurements between groups and conditions, the names of which are stated with the results. Exact p-values are reported where possible, but where tests were performed between conditions on multiple individual animals, p-values are reported as p less than the largest significant p-value in that group after correcting for multiple comparisons, and the number of positive and negative results are stated. Resampling methods are employed in the following specific instances:

For the statistical testing in Figure 5, we tested if the mean turning magnitude in each ΔND or ND bin was different than the bin containing zero, considering each sniff bin separately. This was done by generating a distribution of the differences in resampled means (n=5000) and determining if the confidence interval included zero. Confidence intervals were corrected for multiple comparisons using the Bonferroni method.

To test if nares occlusion introduced left/right biases in following behavior, Figure 6F, we used stationary time-series bootstrapping (Politis and Romano, 1994) to generate simulated distributions of nose position means. The medians of these simulated distributions during individual behavioral epochs were then compared using Wilcoxon rank-sum tests for each animal individually, corrected for multiple comparisons using the Bonferroni method.

## Results

### The Behavioral Arena and Odor Trail Following Task

We constructed a large, open-sided platform on which to perform these behavioral experiments. The size (˜1.3m × 1m) allowed us sufficient space for inter-trial path variability and to obtain optimal behavioral monitoring (see Methods, Figure 1A). We recorded mouse movement by recording high resolution video through a transparent floor, under infrared (IR) illumination via LEDs placed on the edge of the table in an otherwise dark room. This lighting scheme brightly illuminates objects which are near (within a few millimeters) or in contact with the table. We thus obtain high-contrast imaging of the paws, tail, and nose of the mouse (Figure 1B).

**Figure 1.**
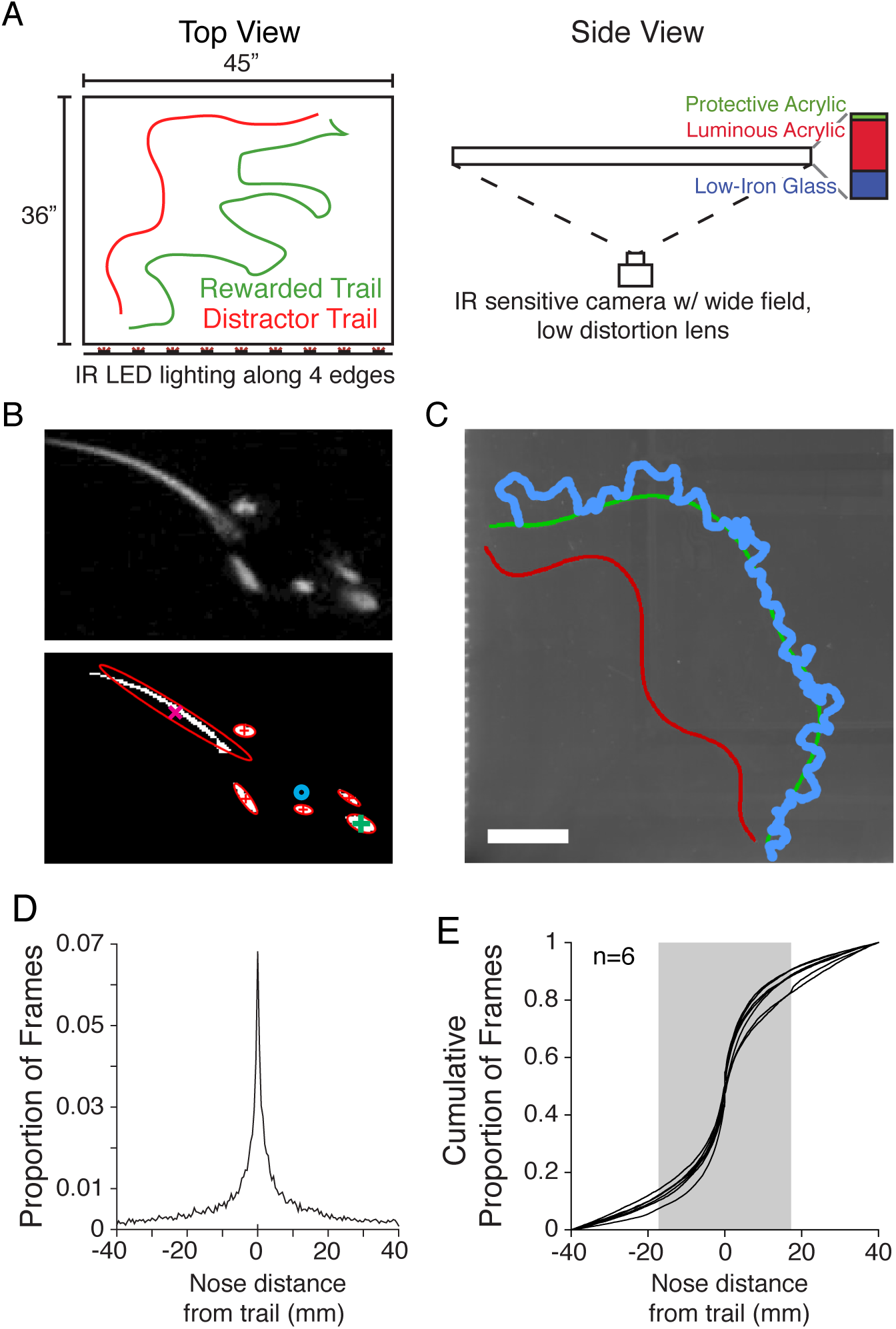
Experimental setup. **A** The large behavioral arena for odor tracking. Odor trails (left), camera position, and the laminate construction of the transparent, luminous floor are shown (right). **B** Example frame crop of showing the mouse after background subtraction (top) and the result of the automated tracking program (bottom). Identified areas are outlined in red, the tail and nose centroids marked with a magenta X and green +, and the mouse center of mass shown with the blue circle. **C** Example nose positions (blue) of a trained mouse during one trial, tracking the rewarded (green) trail. Distractor trail is shown in red. **D** Distribution of nose positions with 40 mm of the trail for one animal. **E** Cumulative distribution of nose positions within 40 mm of the trail for 6 animals during the 3rd week of training.

For each trial, two wax trails (˜4 mm wide), each scented with different low-volatility monomolecular odorants, were drawn by the experimenter onto the surface of the table (Figure 1A, C). We placed small pieces of food at intervals (2-20 cm, depending on the training stage) along the length of the rewarded trail. Trail shape and reward placement along it were varied on each trial so that the animal learned over time that the only reliable predictor of reward placement on the table was the odor trail with the “rewarded” scent. The reward and distractor trail start/end points were each reliably placed near the edge of the table and within approximately 2 cm of each other. Each trial began when the animal was released at one end of the trails, and it was free to explore the table until it had followed the whole trail length, ceased to show interest in the trail for >30 seconds, or 5 minutes had elapsed. Automated tracking of the animal’s nose position (Figure 1B) was then performed on the recorded video (see Methods, Figure 1C).

### Task Learning

After being given two 15-minute sessions to acclimatize to the arena over 2 days, mice were placed in the arena after two short (˜50mm) odor trails were drawn on the floor. One of the trails was heavily baited with food. All food, throughout training, was exclusively placed on the trail with the “rewarded” odorant. Mice quickly learned the basic association, and the training thereafter consisted of gradually lengthening the trails, making them more complex and tortuous, and making the reward placement increasingly sparse. By the end of three weeks of experience, roughly 50 trail exposures over 15 training days, mice would regularly explore continuous sections of trail 400-500 mm in length.

To quantify trail following behavior, we computed the distance from the animal’s nose to the center of the trail (nose distance or ND) on a frame-by-frame basis at typical frame rates of 30-40 Hz. After three weeks of training, the distribution of nose positions relative to the trail is shown in figure 1D for a large area bordering the trail. The animal spends a disproportionate amount of time directly over the trail and much of its time within about 15 mm of the trail. The cumulative density functions of ND for several mice are shown in Figure 1E, and based on these distributions we set a criterion of 17.2 mm for the nose distance from the trail, within which we call “following”. This criterion included 75-80% of a trained animal’s time within 40 mm of the trail, and is wide enough to accept the majority of the movements around the trail the mice would make while following, including casting, where the mouse sweeps its nose back and forth laterally to the trail while walking along it. Typical casting movements are readily observable in the tracking example shown in Figure 1C. Using the above following criterion, we evaluated how quickly animals learn the tracking task by compiling measures of how much of the trail they followed over time. Figure 2A shows the length of trail that mice explored over time for multiple trials and mice. Early in the training, animals mostly followed only small sections, then leave the trail and reacquire it later. This is shown as long horizontal portions of each curve. After training however, mice would often follow the entire length of a trail in 1-2 segments. This improvement, shown as single session snapshots in Figure 2A, is shown throughout training in Figure 2B. Many potential measures of how well the animals perform this task were evaluated. However, we show trail following efficiency because it captures how diligently the animals actually move along the trail. It does not reflect times when the animal was stopped over the trail, and it does not depend on trail length, which increased consistently over the training period. From these data, it is clear that animals start out exploring the two scented trails inefficiently and with only slight preference between them, but learn over time to follow the rewarded one diligently while mostly ignoring the distractor trail.

**Figure 2.**
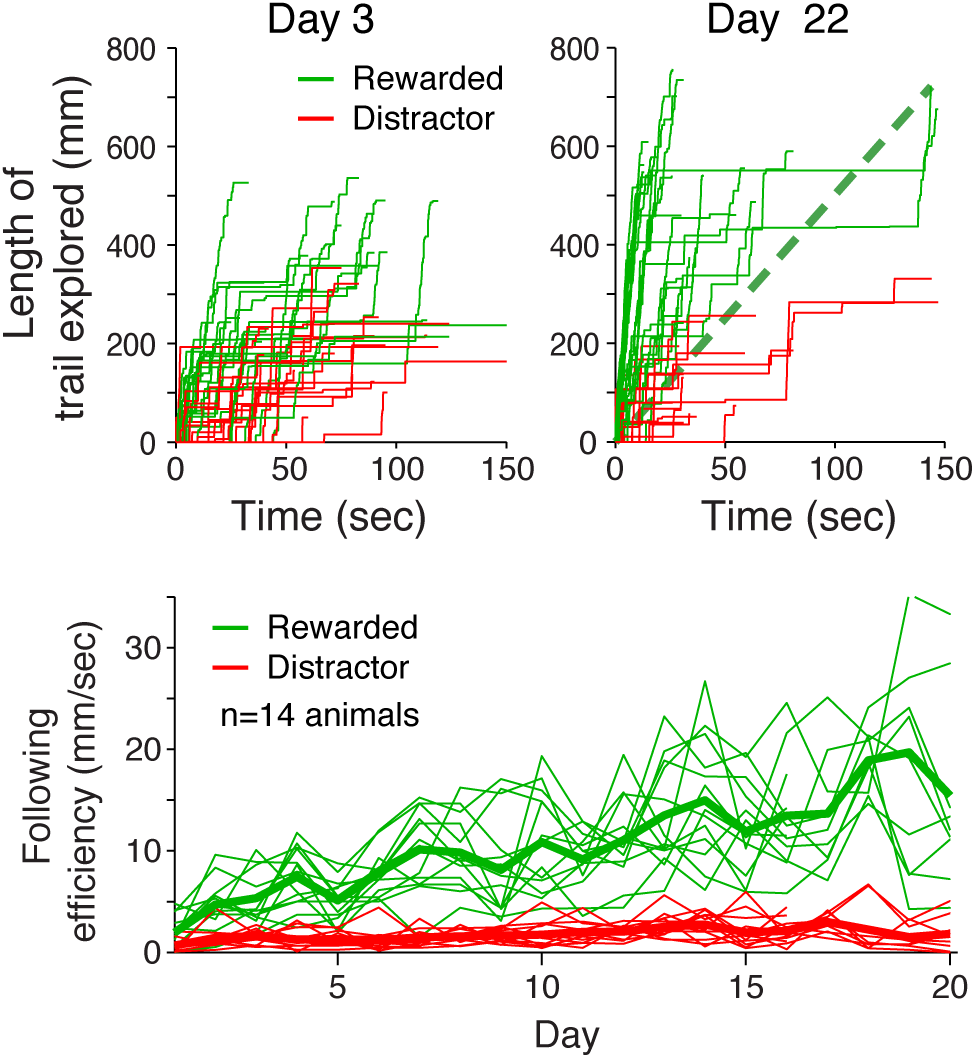
Learning the task. **A** Shows the length of the path explored versus the amount of time elapsed since the trial start, for both the rewarded (green) and distracter (red) trails. In this representation of following, segments of each line with high slope depict times of following and flat segments show times when the animal was either off the trail or stopped along it. Both panels show the performance of multiple mice, during multiple trials, on a single training day. The left shows a day early in training, and the right shows a day after significant experience on the task. **B** Plots the following efficiency metric for each training day. Thin lines show the performance of individual mice, and thick lines show the mean across the population.

### Sniff Patterns during Tracking

We implanted thermocouples into the nasal cavities of a subset of mice trained on this task (n=5) in order to understand how their sensory sampling related to their movements during tracking. All of the animals sniffed at very high rates when following trails, an average of 13.96 Hz (intersniff interval mean/median = 77/70 ms). The high rate of sniffing was punctuated by brief rate decreases that lasted for 1-5 sniffs and were correlated with a decreased nose velocity (Fig 3A). Visual inspection of the video showed some of these events were associated with the animal stopping to eat a food reward. Since such events apparently are not part of active trail following, we excluded from further analysis any data from time periods in which the nose velocity was slower than 50 mm/sec. A significant correlation between the nose velocity and sniff rate remained (p = .035) and the cross-correlogram between of two signals (Figure 3B) indicates that changes in nose velocity lead those of sniff rate by about 40 ms. The relationship between velocity and sniff rate (Figure 3C) shows a strong dependence that saturates at 100 mm/sec. Figure 3D shows that sniff rates across all mice significantly increase when the animal is moving at high velocities (p=10-^8^, Wilcoxon Rank-Sum test). The median sniff rate increase between mouse velocities is 1Hz, indicating that as movement speed increases, the density of trail sampling decreases, despite the increased sniff rate.

**Figure 3.**
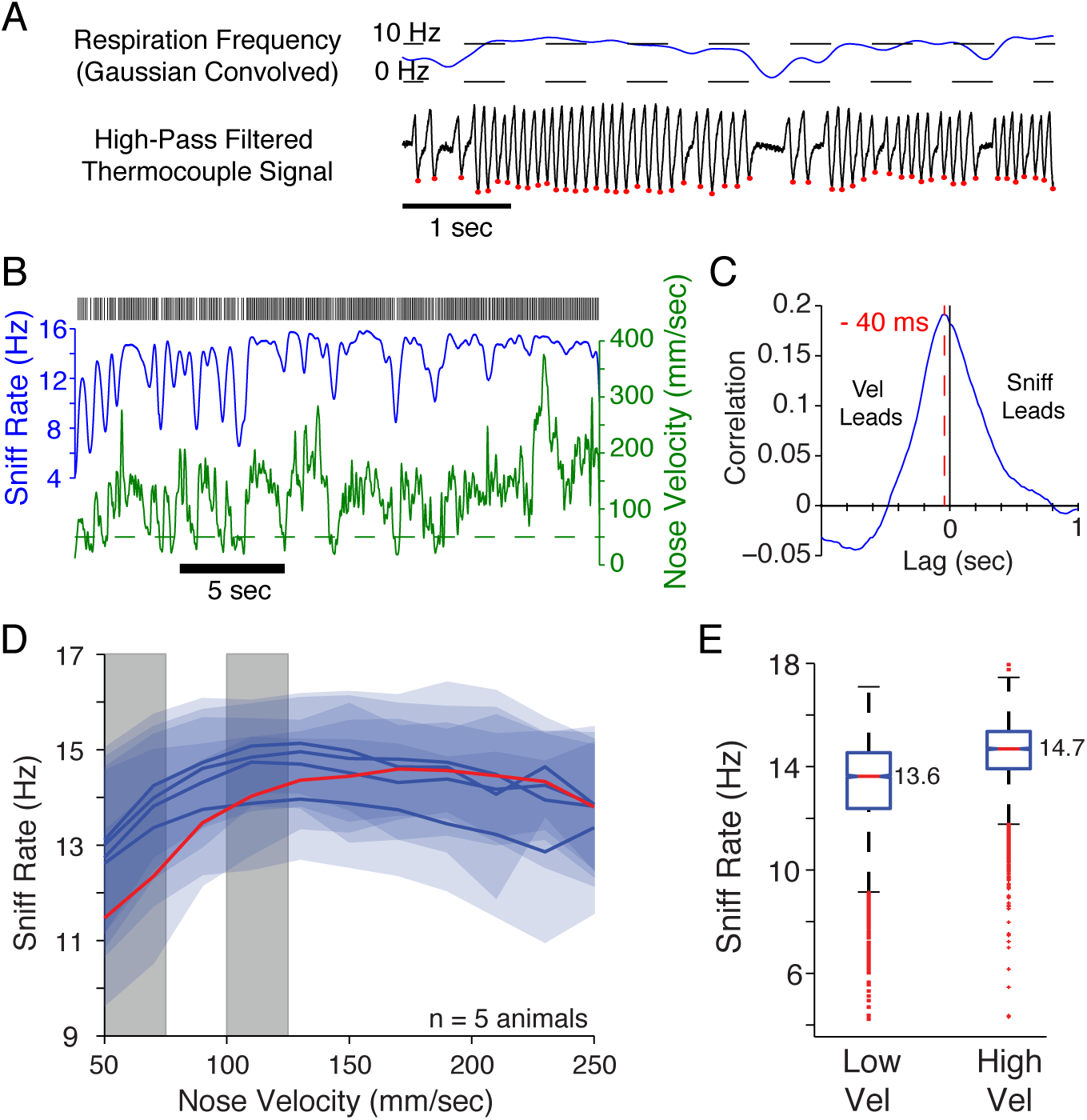
Sniffing during tracking. **A** Example thermocouple output recorded from a mouse while tracking a trail. The black trace shows the thermocouple signal after highpass filtering to remove baseline fluctuations. Inspiration peaks, were then detected via thresholding (red marks), and the resulting binary vector convolved with a Gaussian filter (σ = 50 ms) to obtain a smooth respiration frequency trace (blue). **B** Shows the both the nose velocity (green) and respiration frequency (blue) for a segment of trail following. Black raster on top indicates inspiration times. Dashed line indicates the velocity threshold (50mm/sec) for considering the mouse to be in motion. **C** Cross correlation of the velocity and sniff frequency. The peak correlation occurs when velocity leads the sniff frequency by 40 ms. **D** The average (blue line) and SD (blue shading) sniff frequencies as a function of nose velocity for each of the 5 animals tested. Grey areas mark the velocity ranges used in E. E Shows the sniff rate distributions during low (50-75 mm/sec) and high velocity (100-125 mm/sec) trail following. Box plots indicate the mean, quartile, and span of the distribution with box center, box edges, and whiskers. Red points indicate outliers, and notches indicate 95% confidence intervals on the distribution medians.

### Trail Following Strategy

We next sought to evaluate what sensory cues and strategies the animals employed to following olfactory trails. We limited the mouse to using olfactory cues by eliminating other potential sensory cues. Experiments were performed in a room illuminated only by dim far red filtered light, outside of the mouse visible spectrum. We also recorded several mice (n=7) trained in this task from the side to ensure that they were not touching their noses to the arena floor or visibly whisking the trail. We never observed evidence of a mouse employing these alternative strategies (data not shown). Additionally, the clear preference the mouse displays for the rewarded trail (Fig 2) argues that the mouse is utilizing smell to choose which trail to follow since that is the only distinguishing characteristic of the rewarded trail. Further lateral casting with the head while walking along the trail, a prominent movement pattern that the mouse displays while following the trail, has been observed in the olfactory path following behavior in several other species. This observation suggests that casting is a common strategy employed by a range of animals performing olfactory source localization tasks.

What olfactory cues are the mice then using as they constantly make directional adjustments in order to remain close to the trail? Two available cues are the estimated absolute odor concentration at each sniff and the sniff-to-sniff (intersniff) differences in odor concentration. The former could be compared to an internal expectation of the trail’s odorant concentration in order to estimate distance from the trail while the latter could be used to determine if the animal was moving towards or away from the trail over the time between two sniffs. Intersniff diferences directly provide the change in concentration over samples. A third cue – inter-nostril concentration differences – was considered in later experiments. As a proxy for odor concentration, we measured the distance of the animal’s nose from the trail at the time of each inspiration (ND), and the intersniff change in nose distance from the trail (ΔND). We then examined the animals’ behavior relative to the information provided by each sniff (Figure 4). The probability of the animal making a corrective turn is only minimally influenced by sniff position (ND) for up to 3 sniffs previous (Fig 4B). In contrast, the chance of the animal turning is strongly dependent on the value of ΔND calculated between the most recent two sniffs (Fig 4D). The shape of the relationship between ΔND and turning probability has two peaks – a peak after the mouse has either approached or moved away from the trail. A much shallower dependence is observed when the turning probability is computed using intersniff ΔND from either of the two previous sniff intervals. A large majority of these turns are in the proper direction to keep the animal moving along the trail, with the animal preferentially turning away from the trail for negative ΔND values and turning towards the trail after positive ΔND(Fig 4E). Again, the dependence of the turning direction on ΔNDfor the previous intersniff intervals is much weaker. Similarly, the animal tends to turn towards the trail relative to its position on the sniff immediately previous to the turn (Fig 4C), with weaker and opposite dependencies on the position of the previous two sniffs.

**Figure 4.**
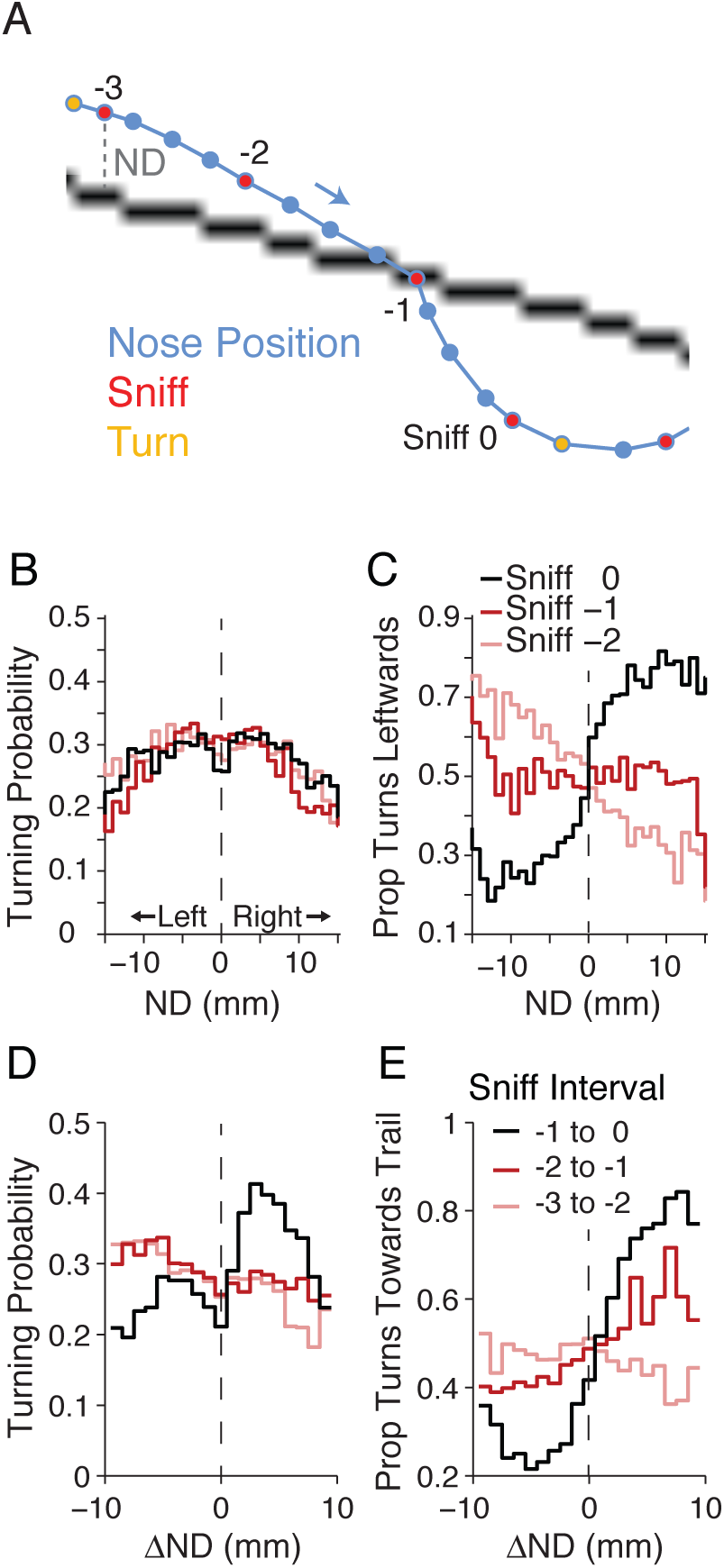
Sniff to sniff determinants of turning behavior. **A** Example of a mouse trajectory with sniffs and turns marked. **B** The probability of turning during the 80 ms interval after a sniff, calculated based on the mouse’s position during each of the 3 preceding sniffs. Same coloring as C. **C** Probability of turning left based on nose distance from the trail center during each of the three preceding sniffs. **D** The probability of turning during the 80 ms interval after a sniff, calculated based on the ΔNDDduring the immediately previous 3 intersniff intervals. Same coloring as E. **E** Probability of the mouse turning towards the trail based on its absolute position during the preceding sniffs.

These data show that after the mouse detects an intersniff concentration change, it is more likely to turn less than 80 ms later (the median intersniff interval). The direction of these turns depends strongly and adaptively on the sign of the concentration change and the side of the trail the animal is on. The effect of absolute distance from the trail on the animal’s decision to turn seems minimal.

Where does this turning strategy take the mouse? To look at this, we segregated the turns into categories based on whether the mouse (or specifically its nose) was approaching the trail (Fig 5 Ai and Aii. ΔND< -2), along the trail (Fig 5 Aiii and Aiv, -2 >ΔND< 2), or away from the trail (Fig 5 Av and Avi, ΔND> 2). We then looked at the movement heading of the mouse relative to the vector pointing towards the trail. Thus, 0° would mean the mouse was moving straight towards the trail, 180° would mean that it was moving directly away, and 90/270° reflect movement parallel to the trail. We observe that the mice often approach and move away from the trail at oblique angles, and their turns also result in movement oblique to the trail (Fig 5A), consistent with the observed casting behavior. The turns they make as they move along the trail seem to result in much more uniform movement directions (Fig 5Aiii-iv). The magnitudes of the turns average about 75-90°, and are significantly larger after the animal moved away from the trail during the previous intersniff interval (Fig 5B; bootstrap, corrected for multiple comparisons). This dependence is not seen for the two previous sniff intervals; in contrast, turns are on average smaller when the animal has moved away during previous sniff intervals (Fig 5C). These data show that the mouse is regularly making corrective turns that take it back and forth at oblique angles along the trail length, and that it mainly uses information about its location change between the two most recent sniffs to modulate the magnitude of those turns.

**Figure 5.**
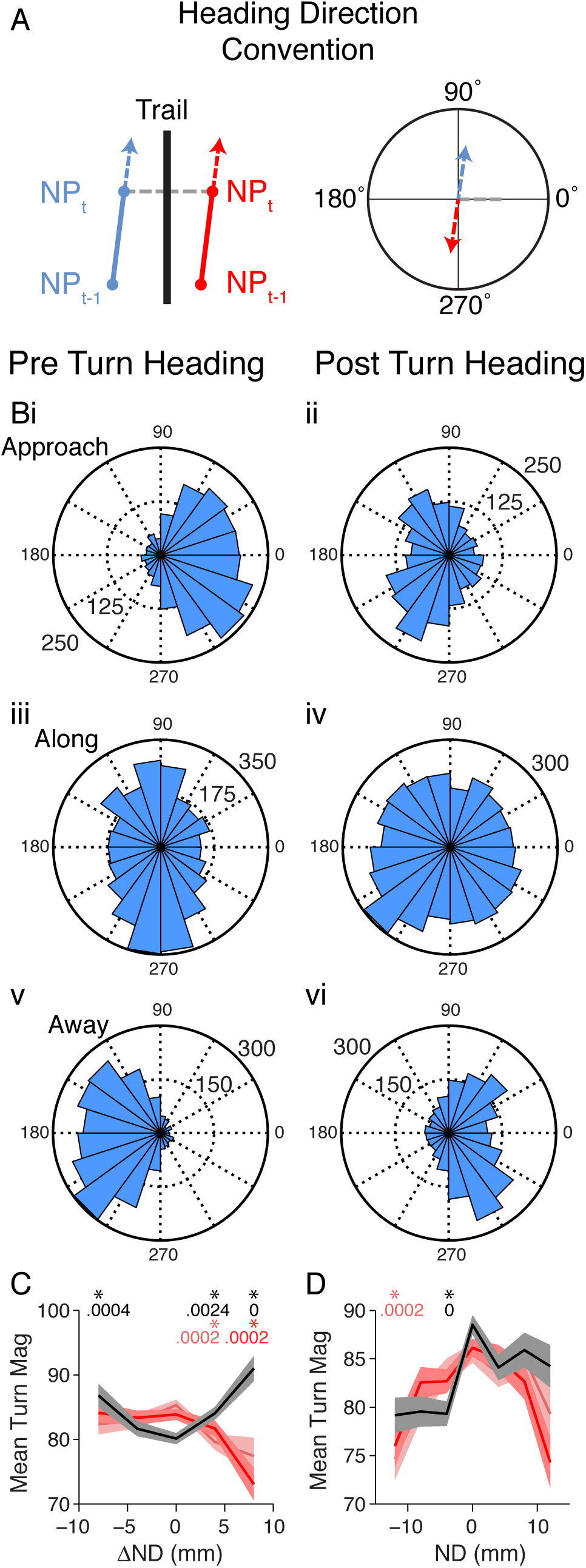
Turning directions and magnitudes depend on the previous sniffs. **A** Schematic showing the direction conventions used. Heading is defined as relative to the vector pointing to the closest point on the skeletonized trail to the mouse’s nose, thus 0 is directly towards the trail, 180deg is directly away, and 90 and 270 deg are along the trail. **B** The pre-turn **(i-iii)** and post-turn **(iv-vi)** headings for when the animal was approaching, moving along, or moving away from the trail. These categories represent ΔNDDvalues of -10 – -2mm, -2-2mm, and 2-10mm respectively during the immediately previous intersniff interval. **C** The mean magnitudes and SEM (lines and shaded areas) of detected turning events, sorted by the ΔNDDof one of the preceding three sniff intervals (same coloring as Fig 4E). Asterisks indicate significant differences in mean turning amplitude from the ΔNDD-2 to 2 bin (p<0.0042, see Methods; uncorrected p values in figure). **D** Turning magnitudes arranged by the absolute sniff position during the previous 3 sniffs (same coloring as Fig. 4C). Asterisks indicate significant differences in mean turning amplitude from the ND bin spanning zero (p<0.0042, see Methods; uncorrected p values in figure).

### Unilateral Nares Occlusion

The use of stereo olfactory cues (i.e. detection of concentration differences between nares) has been proposed as an important cue for olfactory source localization. Indeed, unilateral nares occlusion has been shown to impair navigational performance (Rajan et al., 2006; Porter et al., 2007; Catania, 2013), and has also been shown to effect path following precision in rats (Khan et al., 2012). Therefore, we chose to assess the importance of bilateral cues to mice performing our task using unilateral nares occlusion. We trained animals to follow trails as described above, then reversibly occluded a single nares in each mouse and assessed their behavior for 10-12 days. We then re-assessed behavior after occlusion removal, and again after occlusion of the opposite nares (Figure 6A). We found that occlusion significantly decreased the efficiency with which the mice followed the trail (Figure 6B, see legend for statistics). This reduced efficiency could represent a decrease in motivation to follow the trail or a reduced ability to detect or follow it. In order to distinguish between these two possibilities, and to better understand the behavioral deficit produced by occlusion, we more closely examined the animal’s nose position while following the trail.

If the animal normally uses stereo olfactory cues to localize the trail, we would expect for the animal to systematically mislocalize the trail when one nares is occluded, which would be quantifiable as a bias in the animal’s nose positions relative to the trail when compared to the unoccluded conditions. We observed such bias shifts in the majority of nares occlusions performed, an example of which can be seen in the cumulative distributions of ND for one animal in Figure 6D. Since the animal’s nose position is naturally a time series, with serial correlations between measurements, we treat it as such for further analyses. Plotting the cumulative sum of ND over time (Fig 6E) allows for clearer visualization of how the ND bias changes over time. In this plot, a lateral bias relative to the trail results in a drift away from zero, with negative slopes indicating leftward and positive slopes indicating rightward biases. The animal shown had a slight bias to the left of trail center while unoccluded, but then shifted more leftwards when the left naris was occluded. The bias then shifted back when the occlusion was removed and shifted rightwards when the right nares was occluded. For each animal, we quantified the bias during each experimental epoch (right occluded, left occluded, clear) as the mean position during that time period. Statistical testing of the biases shows that 9 of 10 occlusions resulted in significant shifts in lateral bias (Figure 6F, p< .012 corrected for multiple comparisons, see Methods for details). However, the average magnitude of this shift was small, approximately 1mm in the direction of the nares occluded. We did not observe an effect on the standard deviation around the trail (data not shown), in contrast to observations of trail following in nares occluded rats (Khan et a1, 2013).

**Figure 6.**
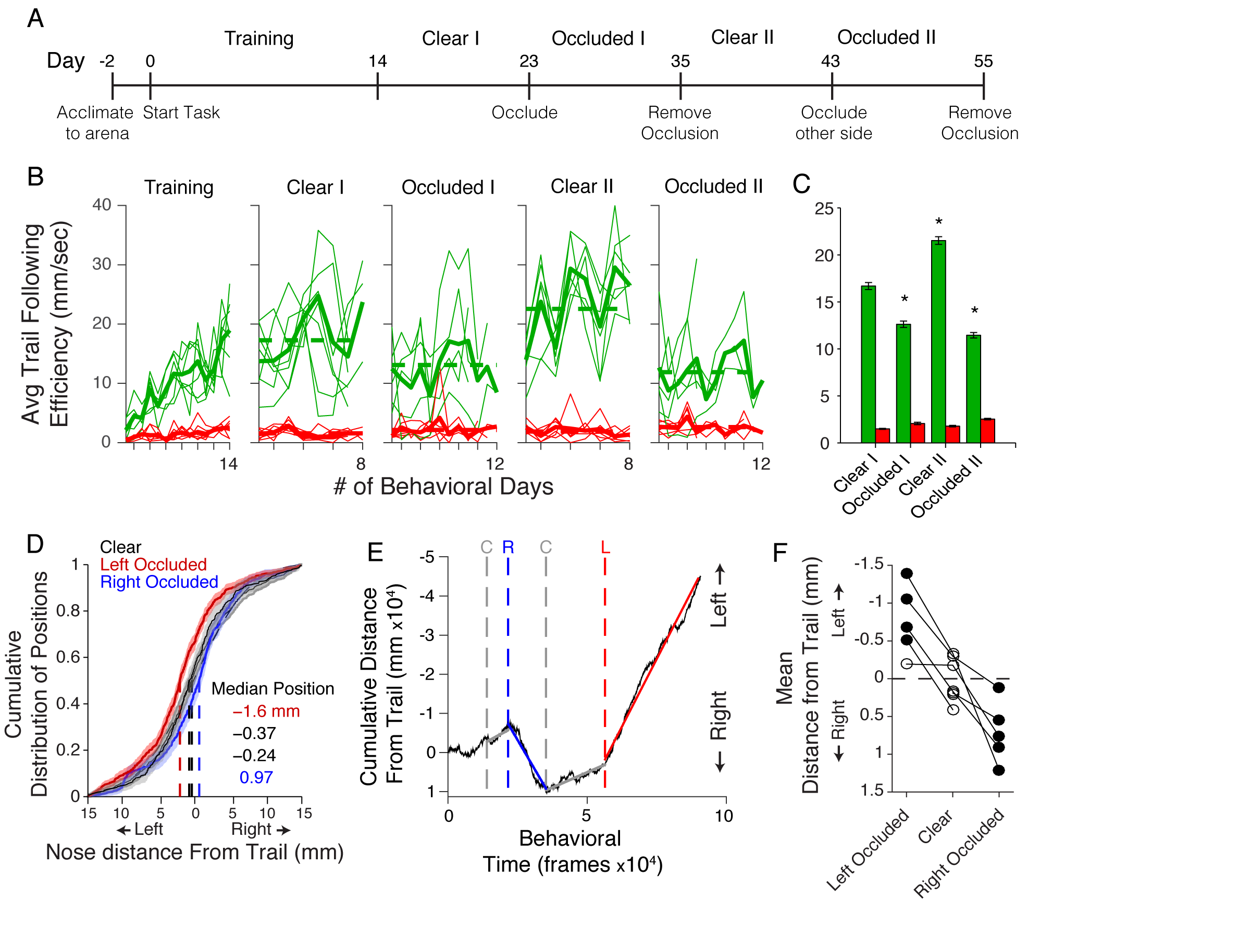
Nares occlusion during trail following biases nose positions **A** The experimental timeline used for training and occluding animals. **B** The following efficiency for individual mice (thin lines) and the average (thick lines) on the rewarded (green) and distracter (red) trails. Dashed lines indicate the mean efficiency during each epoch. **C** Bar graph showing following efficiency during each epoch. Error bars indicate SEM. Stars indicate statistical difference between that condition and the ’clear I’ behavioral period (p =1.5×10^-4^,1.0×10^-5^, 1.6×10^-6^; Wilcoxon Rank Sum). **D** CDF of nose position relative to the skeletonized (1px width) trail for one animal during clear/unoccluded (black), left occluded (red), and right occluded (blue), conditions. Same colors used for panel E. Dashed lines indicate position of the median of each distribution. **E** A cumulative timeseries (black) representation of the nose position of a single mouse, the same mouse as in D. Dashed vertical lines show the start of each experimental epoch. Colored and gray lines show the slopes taken from the mean of the frame-by-frame nose position. **F** Shows the mean shifts in each occlusion condition, for all animals tested. A significant shift from the control condition, pooled over both unoccluded epochs, is indicated by circle fill (p < 0.012, see *Methods* for details).

If nares occlusion were indeed shifting the perceptual position of olfactory trail, one would also expect the animal’s decisions based on the trail’s perceived location to be similarly shifted by occlusion. By casting their noses back and forth while following the trail, animals are effectively making a series of decisions about when the trail is likely in a new direction. Thus, we analyzed the turn positions to see if they were spatially shifted as a result of nares occlusion (Figure 7). We first segregated turns by their direction (rightward/leftward), then averaged them for each behavioral epoch. We find that nares occlusion does indeed shift the positions at which mice turn back towards the trail, and this shift is approximately of the same magnitude and direction as the overall position shift. This observation is consistent with nares occlusion shifting the perceptual location of the trail in the direction of the open nares, implying that stereo olfactory cues are used by mice for precise, millimeter-scale, localization of odor sources.

**Figure 7.**
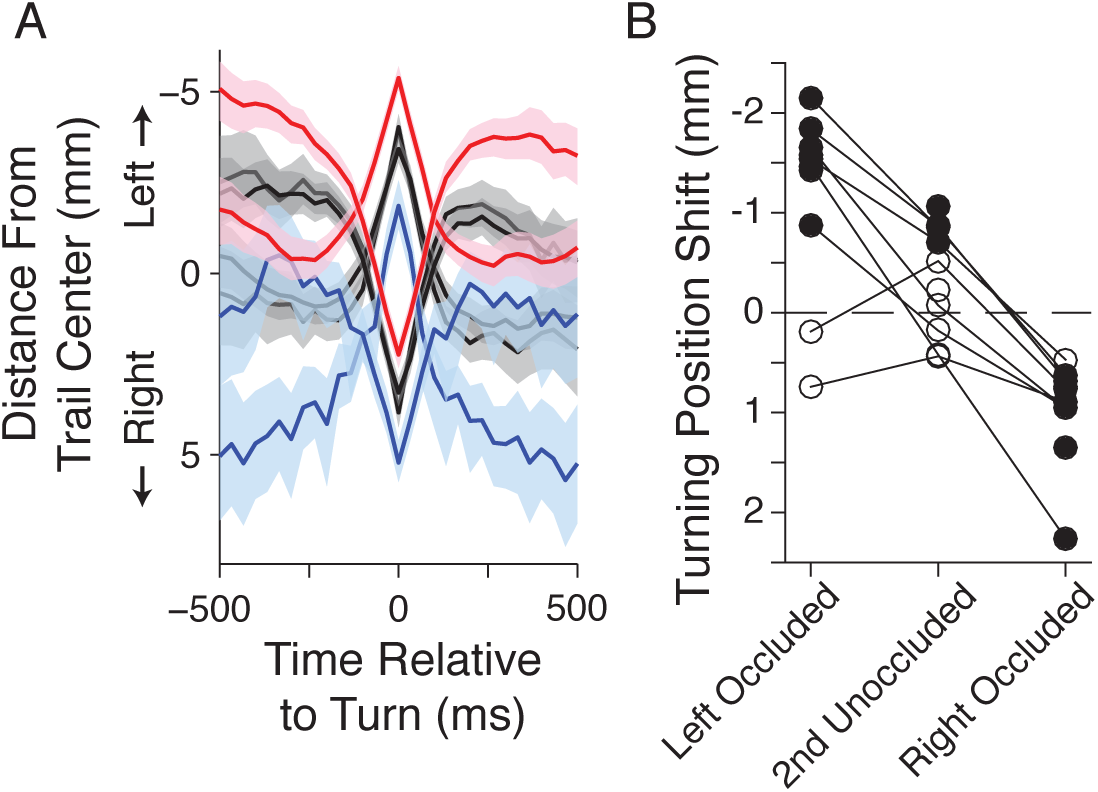
Nares occlusion shifts when animal turning position. **A** Average turn trajectories, separated by the turn direction and by occlusion epoch for an example mouse. Color indicates occlusion condition as in Fig 6D (unoccluded in black, left occluded in red, right occluded in blue) and shaded regions denote SEM of average trajectories. **B** Average turn position shifts relative to turns during the first unoccluded period. Each animal/epoch pair yields 2 shift values, one for each turn direction. Fill indicates a significant shift away from the first unoccluded turning positions (p <0.05, Wilcoxon rank-sum).

## Discussion

We have shown that mice can be trained to follow scented trails that are drawn on a surface, and that they perform this task using a strategy in which they cast back and forth across the trail, with their heads/noses mostly traveling at oblique angles to the trail, making corrective turns back towards the trail whenever they overshoot it. In order to follow the trail efficiently, they employ both a rapid, sniff-by-sniff comparison of odor strength and use stereo olfactory information. We observe that the difference in distance from the trail between the preceding two sniffs is much more predictive of whether or not the animal will turn before the next sniff than the absolute position of the animal or more remote sniff-to-sniff differences in position, suggesting that the difference in intensity between the current sniff and the just previous one is a key cue used by mice to perform this task. The mice react to these sniff by sniff changes in intensity very rapidly, within approximately 80 ms (the intersniff interval observed during this behavior). Furthermore, based on data from unilateral nares occlusion experiments, mice use bilateral sensory information to inform their behavior. Although the way in which this bilateral information is used is not precisely known, occlusion results in a small systematic shift of nose position relative to the trail, suggesting that mice are able to perform some comparison between the intensity of odors arriving at the two nares in a given sniff. This strategy may be most important when the mice are close to the trail where odor concentrations are high and fluctuations may be low. This behavior is most consistent with stereo smelling (i.e comparison of left vs right nares concentration at each sniff), but also may also be consistent with the effect of having two sensors the responses of which are combined in some other way to generate an estimate of distance from the source.

Some previous analysis of odor navigation in rodents has focused on localization of airborne odors (Catania, 2013; Bhattacharyya and Bhalla, 2015; Gire et al., 2016) which may be performed using somewhat different strategies and algorithms than our task. For example, in following airborne odors, determining a consistent wind direction is a key factor in judgments about source localization. Therefore in these scenarios, integrating information about air movement and odor detection will be key to decision making. In our task, although there is no explicit attempt to generate air movement, we believe that ambient air currents are in large part responsible for distributing the odor, since the effective range of molecular diffusion is small over the time intervals that we are studying. Thus, in natural environments with more variable airflow, animal performance level and strategies for following surface based trails could potentially vary with airflow conditions. In previous trail following experiments on rats, performance was in some cases measured only based on latency or success rate (Wallace et al., 2002), whereas in other cases in which nose position was closely monitored, the animals were placed on a moving treadmill, making them unable to stop on the trail or significantly reduce their forward speed and thus to display behaviors that may be important elements of this task (Khan et al., 2012). By employing a large open environment, we have provided maximum behavioral flexibility for the mice and thereby created a task that is unlikely to be solved by relying on non-olfactory cues such as memory, other sensory cues, or a strategy in which the trail is simply occasionally re-discovered as the stimuli shift over time.

Investigations into the neural mechanisms of bilateral olfactory comparisons have focused on the anterior olfactory nucleus (AON) as the likely site of integration for stereo olfactory information. Kikuta et al. (2010 demonstrated the existence of individual neurons with push-pull type responses to ipsilateral and contralateral olfactory stimulation, and recently Rabell et al. (2017) showed that bilateral AONs and the anterior commissure connection between them are both necessary for a directionally appropriate nose shifting response toward a novel odor in head-fixed mice. It is unknown how such behavior observed under head-fixed conditions would translate to orienting behavior in freely moving animals. The neural mechanisms of how sniff to sniff concentration changes may be detected have yet to be explored.

## Acknowledgments

This work was supported by the National Institute of Deafness and Other Communication Disorders-National Institutes of Health (Grant R01-DC011184 to N.N.U.) and the National Science Foundation (NSF 1555916 to N.N.U.). We thank members of the Urban laboratory for helpful comments and discussion.

